# Unraveling the distinct motion bias of TrkA-NGF complex in NGF^R100W^-driven HSAN V disease

**DOI:** 10.1101/2025.02.27.640519

**Authors:** Zahed Bin Rahim, Md Abul Bashar, Nayan Dash, Supti Paul, Rimsha Ziad, Sarmistha Mitra, Raju Dash

**Author notes:** **Corresponding Author Raju Dash** Department of New Biology, Daegu Gyeongbuk Institute of Science and Technology Daegu 42988, Republic of Korea. These authors contributed equally to this study (Zahed Bin Rahim, Md Abul Bashar).

## Abstract

Nerve growth factor (NGF), which binds to tropomyosin-related kinase A (TrkA) receptor, plays essential roles in neuronal survival and function and is also a potent mediator of pain sensation. Mutations in *NGF*, particularly NGF^R100W^, cause hereditary sensory autonomic neuropathy V (HSAN V), which is characterized by insensitivity to pain but without impairment of neurotrophin function. Even though several studies reported the mechanism of growing HSAN V disease, the dynamic mechanisms that dictate its functional specificity remain unclear. In this study, we performed a microsecond scale molecular dynamics (MD) simulation to elucidate the changes in the structural dynamics of NGF by NGF^R100W^ at an atomic level to dissect the distinct motion bias for specific TrkA functions. We found that the NGF^R100W^ reduced NGF dimerization while its binding to the TrkA remained unchanged. NGF^R100W^ enhanced the magnitude of bond formation in the different regions from TrkA, which induced different correlated and dynamic motions associated with impaired nociceptive signaling. The dynamics scenario from this study, shedding light on the deleterious role of NGF^R100W^, provides new structural insights into the function-oriented dynamics motion of the TrkA-NGF complex, offering potential avenues for designing new therapeutics.

## 1 INTRODUCTION

Nerve growth factor (NGF), the first identified member of the neurotrophins family^1^, plays an indispensable role in neurite outgrowth and differentiation, synaptic plasticity, and neuronal survival^1–5^. The gene encoding NGF comprises two promoters that result in two major and two minor mRNA sequences by alternative splicing^6, 7^, where the translation of the two major mRNA transcripts produces preproNGF of 34 kDa and 27 kDa^8^. By eliminating the signal sequence in the endoplasmic reticulum, preproNGF gets converted into 32 and 25 kDa pro-NGF^7, 9–11^, which is, in turn, cleaved by different proteases, resulting in 13.2 kDa mature NGF^12–14^ (**Figure 1a**). It has been known that mature NGF exerts its physiological functions by acting on two types of receptors: the tropomyosin-related kinase A (TrkA) and the low affinity-p75 neurotrophin receptor (p75^NTR^)^5, 15^. NGF exerts its neurotrophic functions by acting on the TrkA receptor^5, 15^. Upon its binding to TrkA, it causes dimerization of receptor and autophosphorylation of distinct tyrosine kinases, thus activating three major downstream signaling cascades, including the PI3K-Akt, The Ras-MAPK-ERK, and the PLC-γ pathways^16–19^, besides the p38 MAPK pathway^20^ and activation of these pathways is responsible for the functions of NGF^18, 21–23^.

**Figure 1.**
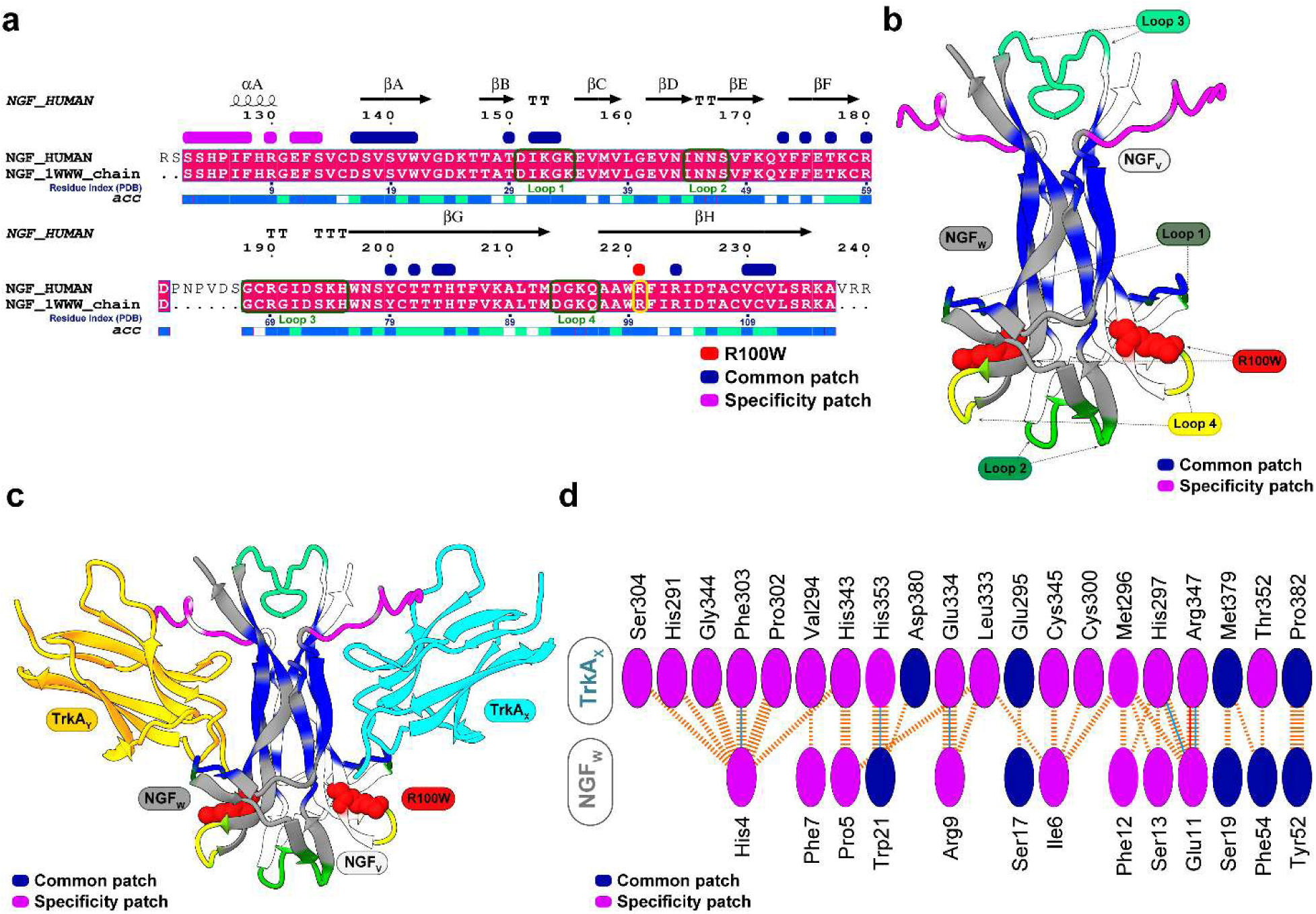
The sequence mapping and structure of NGF with its common and specificity patch, NGF^R100W^ mutation site (221 no. amino acid residue in full sequence), and structure of NGF-TrkA complex with interactions among NGF and TrkA. (a) NGF patch mapping and pairwise sequence alignment of NGF human and NGF 1WWW. The marked regions at the top of the sequence alignment belong to NGF patches, specificity patches (denoted by pink color), and common patches (denoted by blue). The dark green boxes represent the NGF loops (1, 2, 3, and 4), while the yellow box indicates the NGF^R100W^ mutation site (also denoted NGF^R221W^). (b) Structure representing NGF_V_ and NGF_W_ chains and their common and specificity patches and loops. The pink colors denote specificity patches, while the blue color represents the common patch. The green represents the loop regions (1, 2, 3, and 4), and the red shows the NGF^R100W^ mutation site. (c) The structure reveals the binding mode of the NGF-TrkA complex, with light blue representing the TrkA_X_ chain and yellow representing the TrkA_Y_ chain. (d) Inter-chain interactions between NGF chains and TrkA chains were revealed by PDBsum utilizing RCSB PDB ID (1WWW), presenting chains that formed most interactions.

The structure of NGF consists of two sets of twisted antiparallel β -strands, one end of which has three hairpin loops (L1, L2, L4), the other end of which has a reverse turn (L3) and a motif of a cysteine knot (**Figure 1b**). The functional form of NGF forms dimer to exert its physiological function, where two NGF monomers are arranged in parallel and joined by non-covalent interactions^24, 25^. NGF binds to the extracellular domain of TrkA, which is composed of five sub-domains such as (i) two cysteine-rich clusters (domains 1 and 3), (ii) a leucine-rich region (LRR, domain 2), and (iii) two immunoglobulin (Ig)-like domains (domains 4 and 5)^26^. Domain 5 (d5) from TrkA serves as a major site for NGF binding (**Figure 1c**), which is composed of a β-sandwich made up of two four-stranded sheets: one containing strands A, B, E, and D, and the other including strands C′, C, F, and G^25^.Notably, the binding specificity of neurotrophins depends on the specific residues in neurotrophins and Trk receptors, classified as conserved patches and the specific patch. For example, in the NGF-TrkA complex, the conserved patch includes residues that are conserved across all neurotrophins and Trk receptors, while the specific patch pertains to non-conserved residues found at the N-terminus of NGF (residues 2–13) and on the ABED sheet of TrkA-d5^25^, regulating TrkA binding specificity (**Figure 1d**)^25^.

In addition to its nerve growth-promoting role, NGF is a well-recognized modulator of pain pathogenesis with potent pro-inflammatory and sensitizing effects^27–34^. NGF facilitates nociception by binding to TrkA receptors on inflammatory cells, initiating several signaling cascades that increase the production of different inflammatory mediators, for instance, serotonin, histamine, and NGF itself^35^. These mediators are recognized for inducing nociceptor sensitization by altering receptor or ion channel activity at the peripheral terminal^35^. Indeed, several monoclonal antibodies, such as tanezumab^36–38^ and fasinumab^39, 40^, that bind and neutralize NGF effects have shown substantial analgesic benefits above placebo during Phase 2 or Phase 3 osteoarthritis studies.

Accumulating studies have evidenced that mutations in *NGF* are the primary reason for the Hereditary sensory autonomic neuropathy (HSAN) disease, particularly multiple mutations, such as R121W^41^, R221W^42^, and V332fs^43^, in the *NGF* gene were found responsible for HSAN V. R221W, corresponding to R100W (In crystal protein structure) on mature NGF chain, is probably the most studied mutation among them, resulting in a reduction of nociception^42^, fractures of the lower legs and feet^42, 44, 45^, increased cold and heat thresholds^42, 45, 46^, and a decrease in TRPV1, TRPV2, and TRPM8 ion channels responsible for the detection of different thermal stimuli^47^. Interestingly, NGF^R100W^ selectively impaired nociception while maintaining its neurotrophic role intact since the patients with HSAN V do not represent obvious mental impairment but have impaired pain perception^48^. This observation suggests that NGF^R100W^ might trigger specific dynamics motions of NGF that are particularly associated with pain nociceptive signaling but not neurotrophic function.

In this regard, Resende-Lara et al.^49^ identified some collective motions responsible for TrkA activation in the nociception using a normal mode study by comparing wild NGF/TrkA-Ig2 complex with NGF^R100W^. However, serval questions remain about whether NGF^R100W^ can still induce different dynamics motion in NGF in a physiological environment (in the presence of solvent), the effect of NGF^R100W^ in NGF dimerization and NGF/TrkA-Ig2 binding, or responsible conformational changes in NGF or TrkA-Ig2 for different dynamics motion. To answer these questions, this study, therefore, investigates the structural dynamics of NGF/TrkA-Ig2 in the absence or presence of NGF^R100W^ to elucidate conformational and dynamics differences associated with nociception signaling using extensive molecular dynamics (MD) simulation. This study not only uncovers the pathophysiological mechanism for NGF^R100W^ but also explains how distinct functional motion biases NGF signaling.

## 2 RESULTS

To elucidate the structural dynamics of NGF resulting from a mutation at the 100th amino acid, the NGF^Wild^ and NGF^R100W^ were studied by MD simulation at a microsecond timescale. Five unique MD runs of 200 ns each (1 µs in total) were conducted for each condition, and the derived root-mean square deviation (RMSD) analysis for each run in both conditions is shown in **Figure S1**. The RMSD analysis demonstrates that NGF^Wild^ and NGF^R100W^ attained equilibration after approximately 10 ns of simulation and, thereafter, sustained stability. Both systems were found to exhibit a consistent trend across all replicas, indicating their suitability for subsequent analyses.

### 2.1 NGF^R100W^ disrupts NGF dimerization

Before analyzing the effects of NGF^R100W^ on TrkA signaling, it is essential to comprehend the impact of the mutation on NGF dimerization. First, we observed the effect of the NGF^R100W^ on the conformational dynamics of the NGF_V_ and NGF_W_ chains. As shown in **Figure 2a**, NGF^R100W^ reduced the conformation flexibility of the NGF_V_ chain, while a minor reduction was observed in NGF_W_. radius of gyration (Rg) analysis, which indicates protein compactness, demonstrated that mutation enhanced the spatial distribution of its atoms by reducing the compactness of NGF structure (**Figure 2b**), and this change resulted in the dramatic reduction of the solvent-accessible surface area (SASA) of the NGF_W_ chain, suggesting that residues were closely packed in the interior of the protein (**Figure 2c**). However, the change of SASA in the NGF_V_ chain was not substantially noticeable. root-mean square fluctuation (RMSF) analysis, which indicates local residual flexibility, suggested that specific residues from specificity and common patches, as well as loop1 and loop 2, such as Ser17-Trp21, Thr29, Lys34, Asn45-Ser47, and Tyr79 residues, were stabilized in the NGF_V_ chain due to the NGF^R100W^ (**Figure 2d).** A similar trend in the reduction of RMSF was observed for the NGF_W_ chain, where the residues in specificity patches and common patches, such as Ser2-Phe7, Arg9, Glu11, and Ser17-Trp21 residues, retrieved low RMSF (**Figure 2d**).

**Figure 2.**
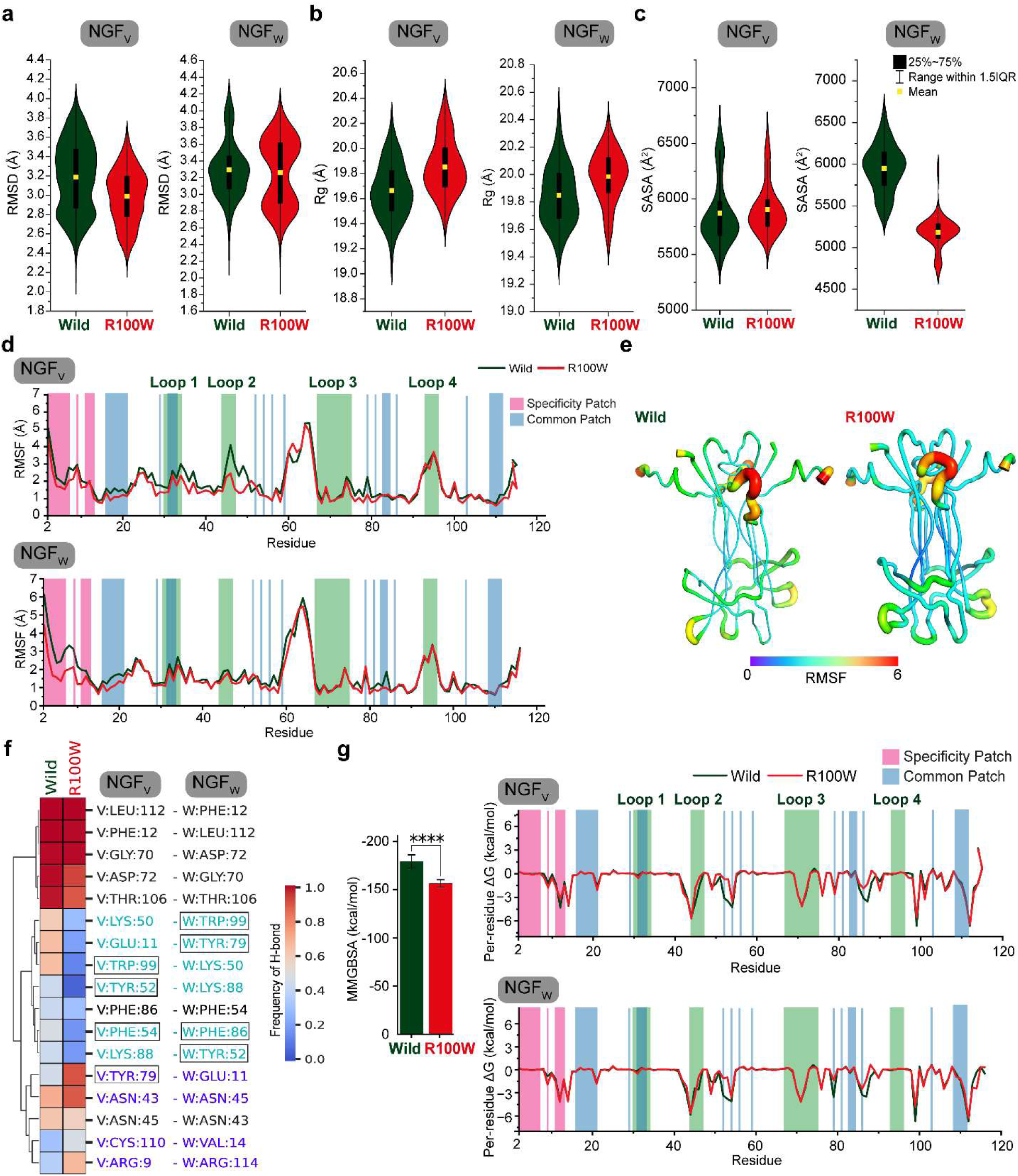
NGF^R100W^ alters the conformational dynamics of NGF and interrupts NGF dimerization. The graphical illustrations represent the mean difference in the NGF conformational alterations between NGF^Wild^ and NGF^R100W^ through (a) RMSD, (b) Rg, and (c) SASA analyses between NGF_V_ and NGF_W_ chains. The residual fluctuations in NGF are displayed based on Cα-root mean square fluctuation (RMSF) through (d) line plot and (e) rainbow color-coded tube representations. (d) The pink and blue colors denote specificity patches and the common patches, respectively, while the green represents the loop regions (1, 2, 3, and 4). (e) A violet-shaded narrow tube represents the region with low RMSF, whereas a red-shaded wide tube denotes the region with high RMSF. The (f) illustration demonstrates interchain H-bond interactions between NGF_V_ and NGF_W_ chains, where the heatmap explains the frequency of H-bond contacts between NGF_V_ and NGF_W_ chains. The aromatic residues with alterations in bond frequency are shown with boxes, and the frequency of bonds that NGF^R100W^ reduces is shown in light blue color, while the frequency of bonds that are raised is depicted in violet color. The (g) graphical representation shows the mean binding energy plot of NGF_V_ and NGF_W_ chains between NGF^Wild^ and NGF^R100W^. (g) The pink and blue colors denote specificity patches and the common patches, respectively, whereas, the green represents the loop regions (1, 2, 3, and 4). The statistical significance between NGF^Wild^ and NGF^R100W^ was examined by utilizing the Student’s t-test: ****p<0.0001. The line plot (H) showing per-residue decomposition analysis shows the binding contribution of NGF_V_ and NGF_W_ chains between NGF^Wild^ and NGF^R100W^.

Since some of these residues deviated in RMSF analysis involved in the dimeric interaction of NGF, we reasoned NGF^R100W^ might alter NGF dimeric interaction. Similarly, the total bond formation between the NGF_V_ and NGF_W_ chains was analyzed using the simulated trajectory, which indicated that NGF^R100W^ reduced the frequency of total bonds to some extent (**Figure S2**). Notably, the most substantial reductions in contact frequency were observed in the residues from the common and specificity patches (**Figure S2**). For instance, Glu11 from the NGF_V_ chain, which had formed contacts with Tyr79 and Val111 of the NGF_W_ chain, was reduced by NGF^R100W^ (**Figure S2**). Likewise, the interaction of Leu112 from NGF_W_, which had established bonds with two residues of NGF_V_, Ser13, and Arg69, was decreased by NGF^R100W^, which also reduced the interactions of Lys50 of NGF_V_ with Lys88 and Trp99 from NGF_W_ (**Figure S2**). In addition, NGF^R100W^ also decreased the interaction of multiple residue pairs between both chains, including Asn45 and Asn46, Tyr52 and Lys88, Thr85 and Phe54, Phe86 and Trp21, and Arg114 and His8 (**Figure S2**), wherein each pair, the former residue belongs to the NGF_V_ chain and while the latter one to that of NGF_W_ chain. In contrast, NGF^R100W^ also increased the contact frequency of some residues, such as the interaction of Arg9 of the NGF_V_ chain with two residues, such as Ser113 and Arg114, from the NGF_W_ chain, was increased by mutation, which also promoted the interaction frequency of His8 of NGF_V_ chain and Arg114 from NGF_W_ chain (**Figure S2**). Moreover, the mutation also increased the magnitude of interaction frequency of several residue pairs between both chains, such as Ser13 and Cys110, Ans43 and Asn45, Leu90 and Ile44, Ala107 and Ala107, Ser113 and Gly10 and Arg114 and Arg9 (**Figure S2**), in which the former residue of each pair comes from the NGF_V_ chain while the latter one from that of the NGF_W_ chain.

Next, we performed hydrogen bond (H-bond) interaction analyses to get more insights, demonstrating a differential perturbation in the H-bond interaction in some residue pairs between the two chains. For example, the magnitude of H-bonding between Glu11 and Tyr79, Lys50 and Trp99, Lys88 and Tyr52, and Phe54 and Phe86 were reciprocal between NGF_V_ and NGF_W_ due to the mutation (**Figure 2f**). However, some residues showed increased H-bond frequency, including the H-bond of Arg9 and Asn43 of NGF_V_ with Arg114 and Asn45 of NGF_W_, respectively, by NGF^R100W^, which also induced the H-bond frequency of Tyr79 and Cys110 of NGF_V_ which had formed H-bonds with Glu11 and Val 14 from NGF_W_ (**Figure 2f**). Nonetheless, the H-bond frequency reduction was more pronounced than the H-bond frequency promotion, indicating that mutation reduced H-bond formation between NGF_V_ and NGF_W_ chains (**Figure 2f**).

As the total bond and H-bond frequencies were reduced due to NGF^R100W^, we hypothesized that NGF^R100W^ might reduce NGF dimer binding energy. Thus, we performed MM-GBSA binding energy analysis (**Table S1** & **Figure 2g**), which showed that NGF^R100W^ decreased the binding energy of NGF dimer from −179.39 ± 6.75 kcal/mol to −156.61 ± 3.81 kcal/mol between NGF_V_ and NGF_W_ chain. Per-residue decomposition analyses indicate that this reduction was mainly in the amino acid residues of the common patch, including Tyr52 and Phe86 residues in both chains, besides Arg9 and Tyr79 in the NGF_W_ chain (**Figure 2h**). These results collectively suggest that NGF^R100W^ disrupts NGF dimer formation.

### 2.2 NGF^R100W^ does not affect TrkA-NGF binding

Since the NGF^R100W^ affects NGF dimerization, we next analyzed whether NGF^R100W^ influences NGF-TrkA binding. As shown in **Figure 3a-b** (left panel), NGF^R100W^ enhances the total number of contacts between NGF and both chains of TrkA. Analyzing the non-bonded interaction frequency revealed that NGF^R100W^ induced differential total bond frequencies of NGF between both TrkA chains, notably in specificity and common patch (**Figure S3**). NGF^R100W^ induced the magnitude of the bonds of several residue pairs between NGF_V_ and TrkA_X_, such as Thr81 and His297, Thr83 and Gln350, Arg103 and Gln350, and Val109 and His297 (**Figure S3**). In addition, the mutation also promoted the frequency of several interactions between NGF_W_ and TrkA_X_ chains, including His4, which formed interactions with Phe303, Gly344, and Cys345. The bond frequencies of Ile6 with Met296 and Cys300 and Pro5 with Glu334 were also increased by NGF^R100W^ in the NGF_W_ and TrkA_X_ chains (**Figure S3**). Similar to the TrkA_X_ chain, the TrkA_Y_ chain experienced an increase in bond frequency. For example, NGF^R100W^ increased the bond magnitude between residues from NGF_V_ and TrkA_Y_, which include Glu11 and His297, Ser19 and Met379, as well as increasing the bond frequency of Trp21 interacting with Thr352 and His353 (**Figure S3**). Furthermore, NGF^R100W^ showed high bond frequency of several residue pairs between NGF_W_ and TrkA_Y_, such as Gly33 and Phe327, Thr83 and Gln350, His84 and Gln350, Phe86 and Gln350, and also the bond frequency of Arg103, which formed interactions with Phe327 and Asn349 (**Figure S3**). However, the magnitude of some residue pairs was also reduced by NGF^R100W^, such as Ser13 and His297 between NGF_W_ and TrkA_X_ chains, Glu11 and Arg347, and Trp21 and Asp380 between NGF_V_ and TrkA_Y_ chains and Val109 and His297 between NGF_W_ and TrkA_Y_ chains.

**Figure 3.**
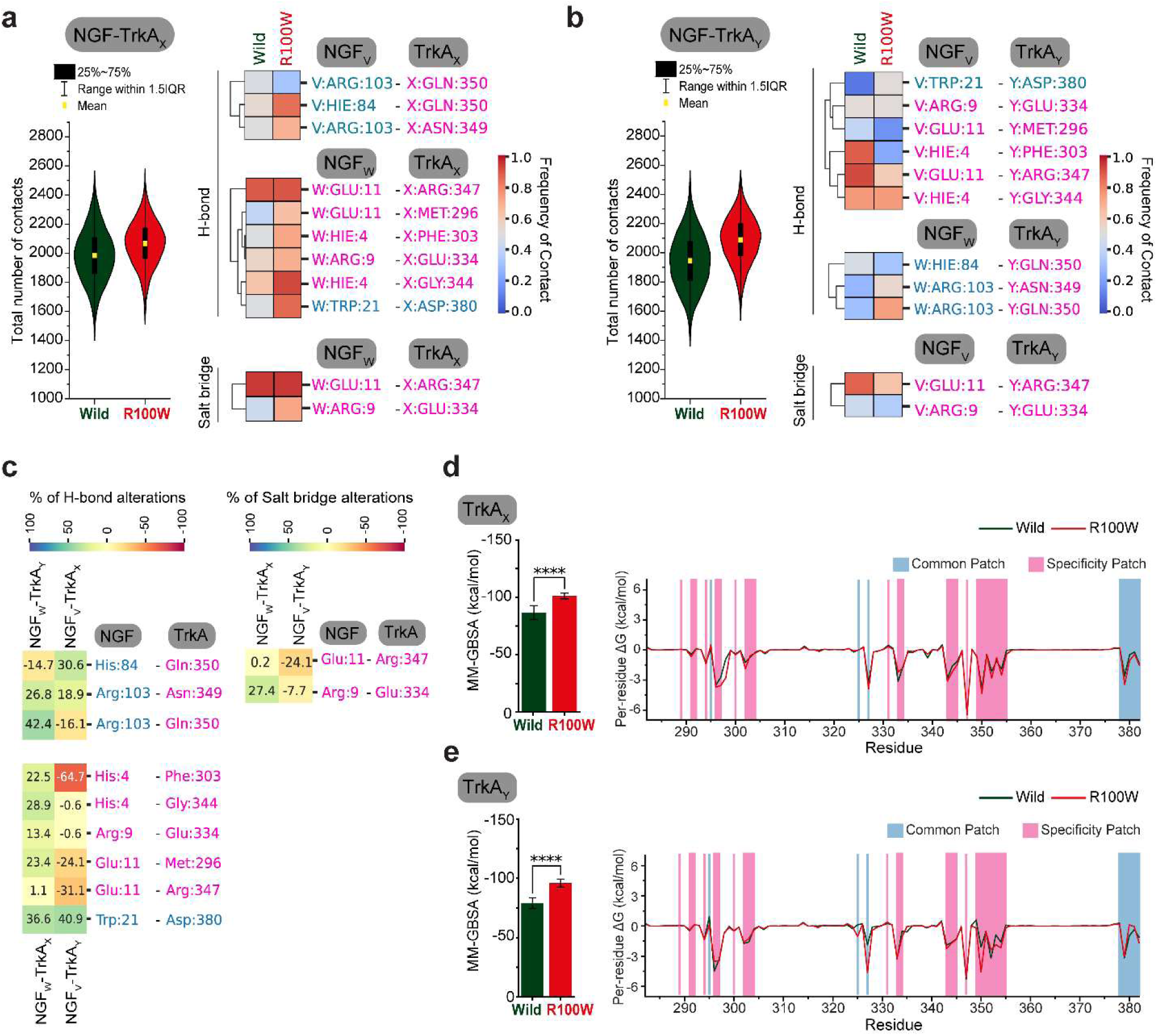
NGF^R100W^ does not affect NGF-TrkA binding. The graphical illustrations present the total number of contacts, H-bonds, and salt bridge frequency formed between NGF and TrkA between NGF^Wild^ and NGF^R100W^, where (a) represents mean total contacts, H-bond and salt bridge frequency between NGF_V_ and TrkA and (b) represents total contacts and H-bond and Salt bridge frequency between NGF_W_ and TrkA. The left panel of (a, b) illustration shows total contact, while the right panel demonstrates the H-bond and Salt bridge frequency. The percentage (%) of H-bond and salt bridge alterations are displayed in illustration (c), where the left panel represents % of H-bond alterations by NGF^R100W^, and the right panel represents % of salt bridge alterations by NGF^R100W^. (a-c) The pink and blue colors denote the residues from the specificity patches and the common patches, respectively. The (d and e) illustrations represent the mean binding energy plot of NGF with TrkA_X_ (d) and TrkA_Y_ (e) on the left panel, and on the right panel (d and e) display per-residue decomposition analysis showing the binding contribution of NGF with (d) TrkA_X_ and (e) TrkA_Y_ chain between NGF^Wild^ and NGF^R100W^. (d-e) The pink and blue colors denote specificity patches and the common patches, respectively. (d-e) Student’s t-test was harnessed to reveal the statistical significance between NGF^Wild^ and NGF^R100W^ in the TrkA_X_ (d) and TrkA_Y_ chain (e): *p<0.05, **p<0.01, ***p<0.001, ****p<0.0001.

Next, we observed that NGF^R100W^ altered the hydrogen bond frequency of several residues from NGF and TrkA (**Figure 3a-b**; right panel). In order to understand these changes in detail, we demonstrated the NGF^R100W^-derived differential changes in H-bond formation in the NGF-TrkA complex using a color-coded heatmap that directly indicates the changes as either a decrease or an increase in bond formation as a percentage (**Figure 3c**; left panel). The heatmap shows apparent changes in differential H-bond formation between different chains of NGF and TrkA complex, especially the residue pairs, such as His84 and Gln350, Arg103 and Gln350, His4 and Phe303, and also, Glu11 which formed H-bonds with Met296 and Arg347 (**Figure 3c**; left panel). Among these residues, drastic changes in H-bonds were observed for the residues in the specificity patch of NGF, particularly His4 and Glu11 (**Figure 3c**; left panel). Indeed, Glu11 showed a reduced salt bridge formation with Arg347 in the case of NGF_v_ and TrkA_Y_ interaction due to the mutation, while the changes in salt bridge formation were not substantial in the case of NGF_w_ and TrkA_X_ (**Figure 3c**; right panel). In addition, Arg9, which is located in the region of the specificity patch, showed differential changes in the case of salt bridge formation (**Figure 3c**; right panel) but not for H-bond, where the h-bond formation of this residue was seen enhanced by mutation with Glu344 in the case of NGF_W_ and TrkA_X_ interaction (**Figure 3c**; left panel). At the same time, a negligible decrease in the salt bridge was observed for NGFv and TrkA_Y_ (**Figure 3c**; right panel). Interestingly, the mutation also increased h-bond formation in all chains for some residue pairs, such as between His84 and Gln350, Arg103 and Asn349, Arg103 and Gln350, and Trp21 and Asp380 (**Figure 3c**; left panel). Nevertheless, collectively, these results revealed that NGF^R100W^ potentiated the magnitude of the bond frequency of NGF with both chains of TrkA more pronouncedly compared to some reductions in the bond frequency, which indicated that NGF^R100W^ induced the total bonds, H-bonds, and salt bridges formation between NGF and its receptor, TrkA, thus promoting their communications (**Figure 3a**-**c**).

Since NGF^R100W^ increased total and H-bonds frequencies, we assumed it may also alter the binding energy between NGF and TrkA. Therefore, we assessed the effect of NGF^R100W^ on binding energy between NGF and TrkA, and the results suggested that NGF^R100W^ induced the MM-GBSA binding energy between NGF and TrkA (**Figure 3d-e**; left panel) (**Table S2**). NGF^R100W^ induced binding energy from −86.63 ± 6.19 kcal/mol to −101.15 ± 2.55 kcal/mol and −78.96 ± 4.54 kcal/mol to −95.90 ± 3.47 kcal/mol in TrkA_X_ and TrkA_Y_ chain, respectively (**Figure 3d**-**e**; left panel) (**Table S2**). The per-residue contribution of binding energy further revealed residues from both specificity and common patches, such as Phe327, Gln350, Thr352, and His353, that contributed to the increase in the binding energy in the TrkA_Y_ chain (**Figure 3d**-**e**; right panel) owing to the mutation. On the contrary, these changes were not significantly pronounced in the TrkA_X_ chain, where His297 mostly contributed to the binding energy extension due to mutation (**Figure 3d**-**e**; right panel). Besides, we also performed per residue contribution for NGF_V_ and NGF_W_ chains forming complex with TrkA (**Figure S4**), which displays that several residues from common and specificity patches contributed to the increase in binding energy contribution between NGF and TrkA, which include residues from NGF_V_ Thr81, Thr83, His84 and residues from NGF_W_ His4, Pro5, Trp21, Arg59 with the TrkA_X_ chain. On the other hand, Arg9, Tyr52, and Arg59 residues of NGF_V_ and Thr83, His84, and Arg103 from NGF_W_ contributed to the increased binding energy with the TrkA_Y_ chain. Overall, these results confer that NGF^R100W^ in NGF does not reduce the NGF-TrkA binding; instead, it induces the frequency of contacts and binding energy between NGF and its receptor, TrkA.

### 2.3 NGF^R100W^ induces conformational alterations of the TrkA

Next, we attempted to discover how NGF^R100W^ affected the various conformational changes of TrkA. As demonstrated in **Figure 4a**, mutation increased the conformational flexibility of the TrkA_X_ chain, which led to a substantial decrease in the compactness (**Figure 4b**) and a slight increase in the SASA of the TrkA_X_ chain (**Figure 4c**), suggesting exposure of hydrophobic amino acids close to the water molecules. However, compared to the TrkA_X_ chain, the effect of the mutation was opposite in the TrkA_Y_ chain, where mutation reduced the conformational changes of the TrkA_Y_ chain, leading to a compact and a minor decrease in the SASA of the TrkA_Y_ chain, indicating a buried structure (**Figure 4c**).

**Figure 4.**
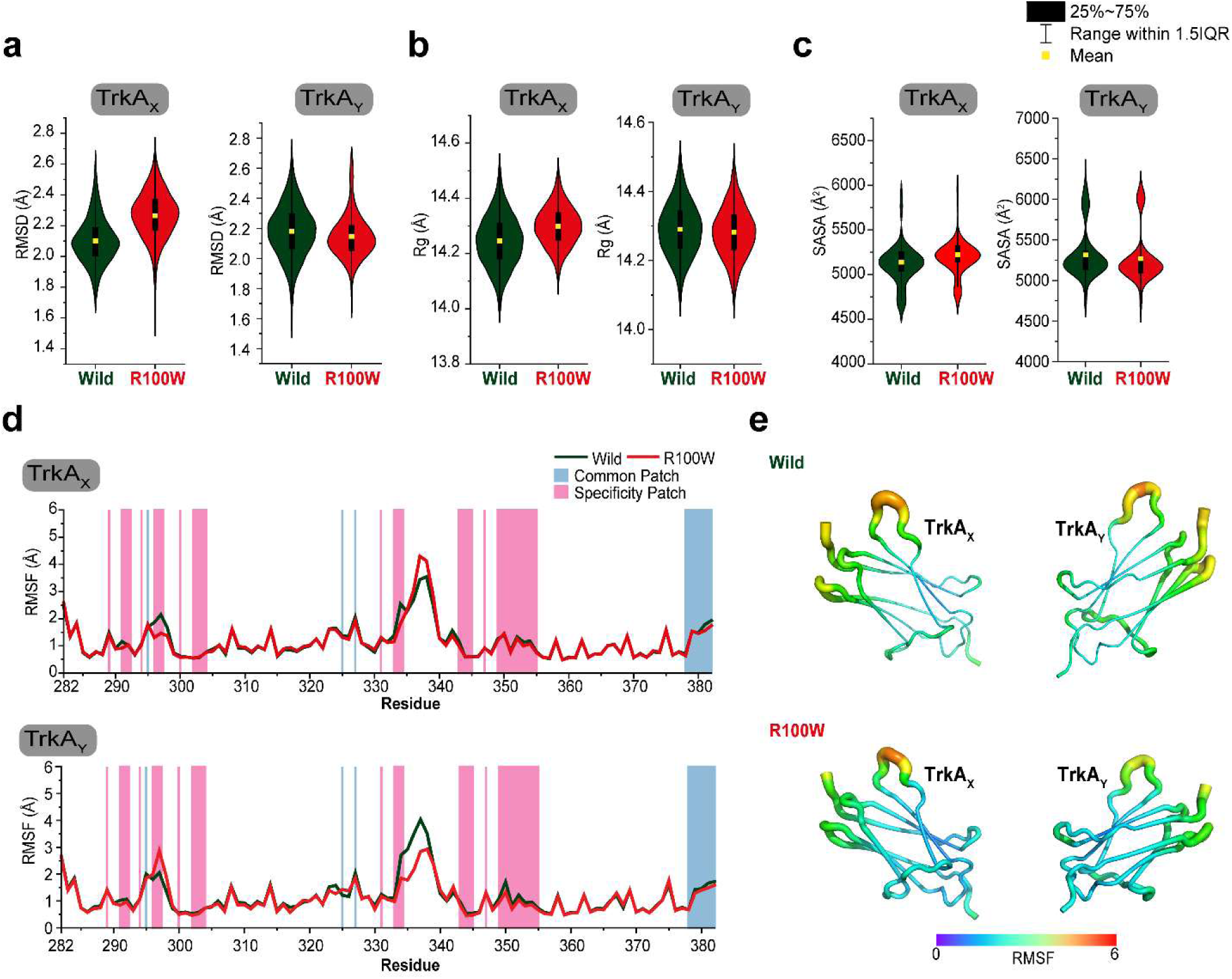
NGF^R100W^ induces conformational alteration of TrkA receptor. The graphical illustrations represent the mean difference in the TrkA conformational alterations between NGF^Wild^ and NGF^R100W^ through (a) RMSD, (b) Rg, and (c) SASA analyses. The residual fluctuations in NGF are demonstrated based on Cα-root mean square fluctuation (RMSF) through (d) line plot and (e) rainbow color-coded tube representations. (d) The colors pink and blue represent specificity patches and common patches, respectively. (e) The region with high RMSF is displayed by a red-shaded wide tube, whereas the region with low RMSF is shown by a violet-shaded narrow tube.

RMSF analysis identified that mutation stabilized two residues (Met296 and His297) from the specificity patch in the TrkA_X_ chain. In contrast, mutation increased the fluctuation of these residues, Met296 and His297, in the TrkA_Y_ chain (**Figure 4d**). Besides, mutation stabilized the Glu334 residue of the specificity patch in both TrkA_X_ and TrkA_Y_ chains (**Figure 4d**). These results, taken together, suggest that the NGF^R100W^ promotes different conformational dynamics of TrkA.

### 2.4 NGF^R100W^ changes the dynamic motions of the TrkA

Since NGF^R100W^ induces different conformational changes in TrkA, we sought to analyze whether it can alter the dynamic motions of TrkA. Thus, we first investigated the effect of NGF^R100W^ on inter- and intra-residual correlated motion of both chains of TrkA. As shown in **Figure S5**, mutation significantly altered the correlated and anti-correlated motion in both chains of TrkA, and these alterations differed in both chains. NGF^R100W^ induced both correlated and anti-correlated motion in the TrkA_X_ chain, but in the TrkA_Y_ chain, it reduced these motions to random (**Figure S5**). In order to identify the regions which experienced substantial alterations in motion due to NGF^R100W^, we further plotted the absolute differences of dynamics cross-correlation matrix (DCCM) of TrkA_X_ and TrkA_Y_ (based on **Figure S5**) between NGF^Wild^ and NGF^R100W^ in **Figure 5a** (both left and right panels), which depicted mutation-induced correlated and anti-correlated motions of several regions (marked by a square bracket in blue color) in the TrkA_X_ chain, such as Thr292-Pro302, Thr330-Asn356, and Phe378-Pro382, where most of these residues were located in the common and specificity patches (**Figure S5a** & **Figure 5a**). However, compared to the TrkA_X_ chain, the differences in DCCM were less pronounced in the TrkA_Y_ chain, where NGF^R100W^ reduced correlated and anti-correlated motions to random motions as observed in His297-Gln308 and Phe332-Arg347 regions (**Figure S5b** & **Figure 5a**).

**Figure 5.**
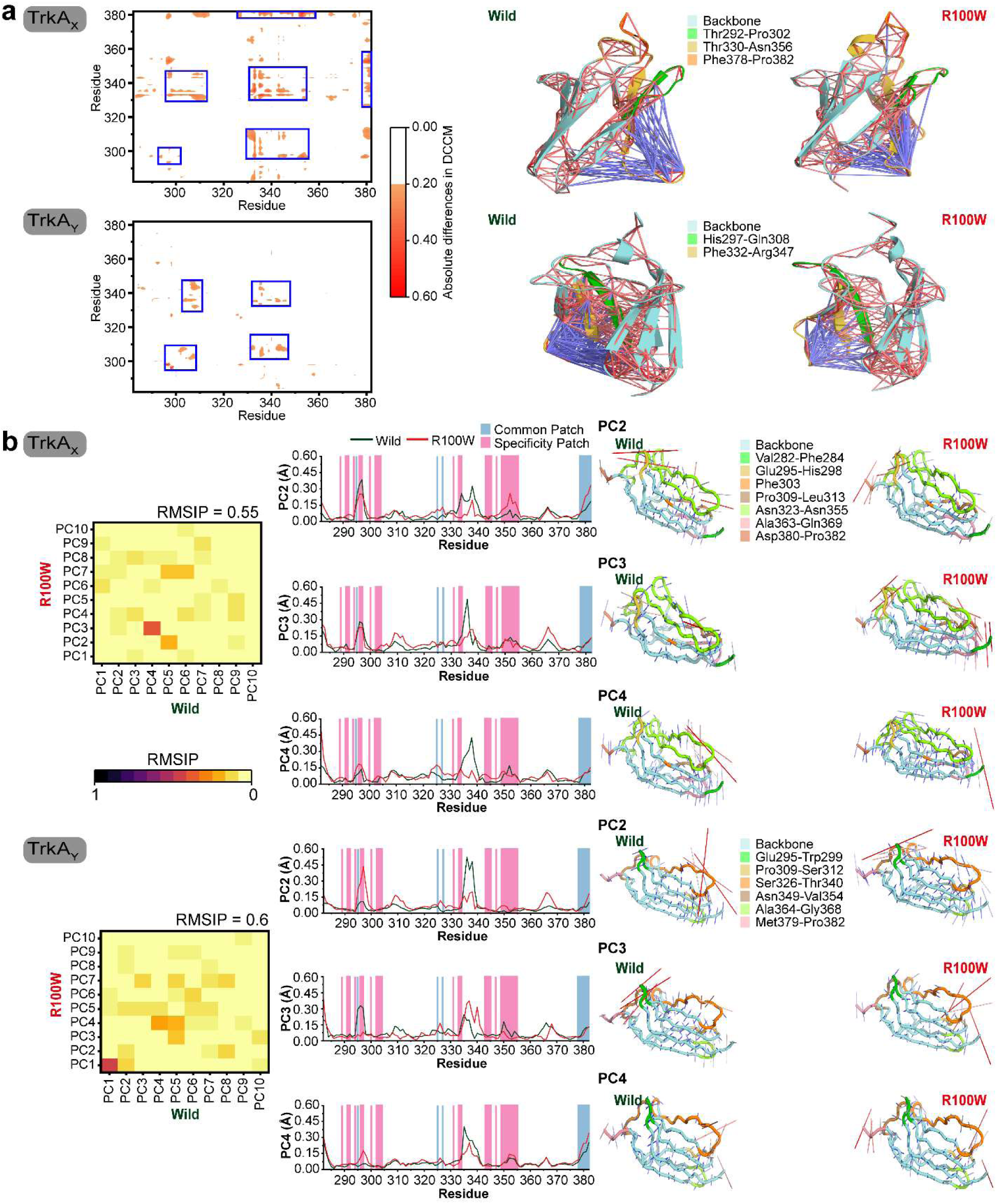
NGF^R100W^ displays alterations in correlated and anti-correlated motions as well as alterations in principal component analysis. (a) The illustration (left panel) shows the absolute differences in the dynamic cross-correlation matrix (DCCM) between NGF^Wild^ and NGF^R100W^, where a DCCM difference of more than 0.2 is represented in orange colors. The upper plot represents the absolute difference in the DCCM plot for the TrkA_X_ chain, while the bottom plot shows the absolute difference in the DCCM plot for the TrkA_Y_ chain. The illustration (right panel) represents the structural view of DCCM, where blue lines (−4 to −10) indicate a negative correlation and red lines (4 to 10) indicate a significant positive correlation. The degree of strong correlation is apparent through the intensity of the line colors. The variations in DCCM of different regions from TrkA_X_ and TrkA_Y_ chains are displayed in green, yellow, and orange, with the backbone structure in cyan color. (b) To illustrate the similarities and variations in the conformational spaces of the NGF^Wild^ and NGF^R100W^, including structures, the root-mean-square inner product (RMSIP) values of the first 10 PCs were calculated and depicted as a gradient heat map (left panel) ranging from yellow to dark red to signify a low and a high value. The upper gradient heat map displays the first 10 PCs for the TrkA_X_ chain, while the bottom heat map shows the first 10 PCs for the TrkA_Y_ chain. The middle panel (b) displays line plots illustrating the mobility degree recorded in PC2, PC3, and PC4 for the TrkA_X_ chain (upper plots) and TrkA_Y_ chain (bottom plots) due to NGF^Wild^ and NGF^R100W^, where the specificity patches and common patches are indicated by the colors pink and blue, respectively. The right panel (b) displays the contributions of PC2, PC3, and PC4 to NGF^Wild^ and NGF^R100W^ using porcupine plots, highlighting the variations in magnitude and direction of motions by the length and direction of mod vectors in the TrkA_X_ (upper plots) and TrkA_Y_ (bottom plots) chain. The variations in motions of different regions from TrkA_X_ and TrkA_Y_ chains are displayed in distinct colors, ranging from red to green and yellow, with the backbone structure in cyan color.

Next, in order to explain the dominant movement by overall conformations that differ between the NGF^Wild^ and NGF^R100W^, we conducted principal component analysis (PCA), which reveals the collective motions of the localized fluctuations with the help of PCs. Similar to DCCM analysis, the pairwise comparison of the top 10 PCs between NGF^Wild^ and NGF^R100W^ utilizing RMSIP calculations pointed out that dynamic motions were distinguishable between both states, NGF^Wild^ and NGF^R100W^ (**Figure 5b**; left panel), where most of the protein conformational variations were recorded by PC1, succeeded by PC2 and PC3 (**Figure S6**). Remarkably, PC1 of TrkA-NGF^R100W^ captured a higher proportion of dynamics of TrkA_x_ than PC1 of TrkA-NGF^wild^ (**Figure S6a**) but surprisingly did not display any alteration in the TrkA_x_ dynamic motion, and it was also true for TrkA_Y_ (**Figure S7**). On the other hand, PC5 showed a negligible change between TrkA-NGF^wild^ and TrkA-NGF^R100W^ (**Figure S7**), whereas PC2, PC3, and PC4 showed notable fluctuations in both chains, especially in several regions, such as Val282-Phe284, Glu295-His298, Phe303, Pro309-Leu313, Asn323-Asn355, Ala363-Gln369, Asp380-Pro382 in the TrkA_X_ chain, and Glu295-Trp299, Pro309-Ser312, Ser326-Thr340, Asn349-Val354, Ala364-Gly368, Met379-Pro382 in the TrkA_Y_ chain and most of these residues were from specificity and common patches (**Figure 5b**; middle panel). Since PC2, PC3, and PC4 captured substantial fluctuations, they were additionally illustrated in porcupine plots to observe the direction of structural shifts. According to **Figure 5b** (right panel), NGF^R100W^ altered the direction of structural shifts of the regions (colored regions: red, green, and yellow) captured by PC2, PC3, and PC4 in both TrkA_X_ and TrkA_Y_ chains. These observations, obtained from DCCM and PCA analyses, collectively indicate that NGF^R100W^ alters the conformational ensembles of the TrkA across several important regions, which include Glu295-His298, Thr330-Asn355, Asp380-Pro382 regions in the TrkA_X_ chain and His297-Gln308, Phe332-Arg347 regions in the TrkA_Y_ chain.

## 3 DISCUSSION

Due to the intrinsically dynamic nature, each protein may confer different conformations in solution, and these differential conformations are substantial for its particular biological functions^50, 51^ and enable each protein to exert multiple functions^52, 53^. Hence, the functions of proteins depend on their particular folding and, eventually, their dynamics. Mutations in proteins may alter their conformations, dynamics, and stability^54^, which may abolish the functions of proteins and/or gain new ones. This way, mutation may exert a critical role in pathophysiology, and understanding the change of protein conformation and dynamics due to the mutation not only helps to understand the disease mechanism but also explains how specific motion in a protein determines a particular function. In this study, we performed MD simulation at a microsecond scale to elucidate how NGF^R100W^ altered conformational dynamics of NGF and eventually associated dynamics motion to suppress the NGF-TrkA-triggered nociceptive signaling but not neuronal growth^42, 48^.

In this study, we identified that NGF^R100W^ induces the differential conformational dynamics (**Figure 4a-c**) in an opposing pattern for the two chains of TrkA, such as stabilizing Met296 and His297 in the TrkA_X_ chain but increasing their residual flexibility in the TrkA_Y_ chain (**Figure 4d**). In contrast, the mutation stabilized Glu334 in both the TrkA_X_ and TrkA_Y_ chains (**Figure 4d**). Interestingly, these residues are located in the key regions having altered correlated and anti-correlated dynamics motion in TrkA-NGF^R100W^, specifically, Glu295-His298, Thr330-Asn355, Asp380-Pro382 regions in the TrkA_X_ chain and His297-Gln308, Phe332-Arg347 regions in the TrkA_Y_ chain, according to the DCCM and PCA analyses (**Figure S5** & **Figure 5**). We suggest that dynamics flexibility in these regions is important for preserving TrkA-mediated nociceptive signaling, and alteration might be associated with signaling disruption. Notably, the substantial difference in dynamic motion between wild and mutant was only observed in the case of PC2 to PC4 but not in PC1, which covered the maximum proportion of variance in dynamics for all systems (**Figure S6-7** & **Figure 5b**). We suggest that this motion involved in PC1 is associated with neurotrophin signaling, which explains why NGF^R100W^ is ineffective in disrupting neurotrophic function.

However, it is also crucial to understand how NGF regulates TrkA dynamics motion in the TrkA-NGF^R100W^ complex. Analyzing intermolecular interactions of the NGF-TrkA complex in both wild and mutant suggests that several residues in NGF^R100W^, such as His4-Ile6, Arg9, Glu11, Ser13, Ser19, Trp21, Gly33, Thr81, His84, Phe86, Arg103, and Val109, which maintain key interactions with the several residues of TrkA, such as Met296, Cys300, Phe303, Phe327, Glu334, Gly344, Cys345, Arg347, Asn349, Gln350, Thr352, His353, Met379, and Asp380, act as major regulators for altered TrkA motion in TrkA-NGF^R100W^ (**Figure S3** & **Figure 3a-c**). Indeed, these residues are located in the dynamically influenced regions of TrkA identified by DCCM and PCA analyses (**Figure S5** & **Figure 5**), and their key interactions with NGF have been suggested as substantial for TrkA activation^25, 56^.

Our study also revealed that NGF^R100W^ does not reduce the binding energy between NGF and TrkA (**Table S2** & **Figure 3d-e**), indicating that it does not disrupt the NGF-TrkA complex, which is consistent with a previous study in which Covaceuszach et al.^55^ characterized NGF-TrkA binding using Surface Plasmon Resonance. However, we observed that NGF^R100W^ reduced the NGF dimer formation. First, NGF^R100W^ induced altered structural dynamics of NGF (**Figure 2a-d**) and reduced interactions of several aromatic amino acid residues (**Figure S2** & **Figure 2f**), including His8, Trp21, Tyr52, Phe54, Tyr79, Phe86, and Trp99, which were reported to be essential for NGF dimer stability previously^57, 58^. Furthermore, NGF^R100W^ reduced dimer binding energy in MM-GBSA analysis, where Arg9 and some aromatic residues such as Tyr52, Tyr79, and Phe86 contributed to the binding energy reduction (**Table S1** & **Figure 2g**).

Collectively, this study concludes that NGF^R100W^ reduced NGF dimerization and induced several intermolecular interactions between NGF and several distinct regions from TrkA, including residues from Glu295-His298, Thr330-Asn355, Asp380-Pro382 regions in the TrkA_X_ chain and His297-Gln308, Phe332-Arg347 regions in the TrkA_Y_ chain, leading to differential conformational dynamics and altering directions of motions of TrkA, and thus contributing the pathological consequences of NGF^R100W^ causing HSAN V disorder.

## 4 MATERIALS AND METHODS

### 4.1 Preparation of the simulation system and molecular dynamics (MD) simulations

The crystal structure of the NGF was downloaded from the RCSB protein databank with the bearing PDB ID of 1WWW. Based on the previously described procedure^59–64^, this structure was then prepared utilizing the Schrödinger 2023-2 (Schrödinger, LLC, New York, NY, USA). Following this, the same software was utilized to generate the mutant, NGF^R100W^, with the help of residue scanning calculation within Schrödinger 2023-2^59, 65, 66^.

To ascertain the conformational changes in the NGF structure induced by mutation, we conducted MD simulation by harnessing the academic edition version of the Desmond program Schrödinger 2023-2 (Schrödinger, LLC, New York, NY, USA)^67, 68^. The structures were modeled by employing OPLS4 force field and were positioned inside an orthorhombic box with a dimension of 10 Å in every direction, based on previously described methods^61, 69–72^. Utilizing the explicit solvation model, that is, the Monte Carlo simulated transferable intermolecular potential 3 points (TIP3P) water model, we solvated the NGF^Wild^ and NGF^R100W^ structure. In order to achieve neutralization, we added counterions (Na+/Cl^-^), and the system’s salt content was altered to 0.15 M to imitate the physiological environments. Prior to conducting the MD simulation, we minimized and equilibrated each structure by harnessing the default Desmond protocol, which consists of a series of constrained minimization phases and MD based on the previous report^73^. The MD simulation was performed under thermodynamic conditions utilizing the isotropic Martyna-Tobias-Klein barostat ^74^ and the Nose-Hoover thermostat^75^ to operate pressure at 1 atm and temperature at 300 K. The long-range electrostatic interactions were analyzed employing the particle mesh

Ewald method, and the short-range electrostatic interactions were evaluated with a cut-off of 9.0 Å. Within a preset short-range cut-off, the multistep RESPA integrator was employed to integrate the equations of motion for both bonded and non-bonded dynamics, and this integration was performed with a 2.0 fs inner time step^76^. An outer time step of 6.0 femtoseconds accounted for non-bonded interactions that exceeded the cut-off. Finally, utilizing the NPT ensemble approach, the MD simulation was run 5 times independently for each structure (NGF^Wild^ and NGF^R100W^), with each simulation performed for a duration of 200 nanoseconds, and the coordinates were recorded after every 100 picoseconds. Subsequently, we monitored the stability of NGF^Wild^ and NGF^R100W^ utilizing the trajectories of MD simulation through RMSD, RMSF, Rg, and SASA via Schrödinger 2023-2.

### 4.2 Analysis of protein dynamics

The DCCM was carried out using an R package Bio3D to understand motions that correlated to time by amino acid residues throughout the MD simulation^77^. The equation harnessed to assess the cross-correlation ratio (C_ij_) between the skeleton carbon atoms i and j is as follows.

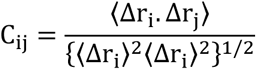

Here, Δr_i_ and Δr_j_, respectively, stand for the average location of the ith and jth residues, while the angled brackets “〈〉” denote the mean time of the whole trajectory. The DCCM values demonstrated a range from −1 to +1, with positive values indicating a positively correlated motion and negative values indicating a negatively correlated motion.

Additionally, the flexibility and collective motions of NGF^Wild^ and NGF^R100W^ were explicated by PCA, which was computed with the help of an R package Bio3D. After eliminating translational or rotational components, we examined both the covariance matrix of atomic coordinates and the eigenvectors in order to get the PCA eigenvectors values. Following this, the orthogonal coordinate transformation matrix underwent diagonalization, producing the diagonal matrix of eigenvalues. The columns represented the eigenvectors that correlated with the direction of motion measured in relation to the beginning coordinates. The eigenvector was linked with the eigenvalue and denoted the system’s total mean-square displacements (MSD). Previous works have mathematically addressed further details on the entire system^78, 79^.

### 4.3 Trajectory clustering analysis

We utilized Desmond’s trajectory clustering analysis tool to group the complex structure retrieved from the comprehensive MD simulation trajectory by employing the backbone RMSD matrix of the NGF-NGF and NGF-TrkA complexes. Clustering was performed in a maximum of 10 clusters with a frequency of 5. Each cluster then underwent MM-GBSA binding energy calculation via a web server, HawkDock^80^, and the specifications were previously detailed in^81^.

### 4.4 Percentage of bond alteration analysis

For better understanding, the interaction frequency of bonds (H-bonds and salt bridges) between NGF^R100W^ and NGF^Wild^ obtained from contact analysis were subtracted, and then the subtracted value was multiplied by 100 to get the percentage of bond alteration by NGF^R100W^. The following equation was harnessed to get the percentage of bond alterations.

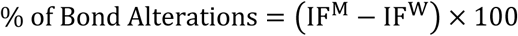

Here, IF^M^ and IF^W^ indicate the interaction frequency of NGF^R100W^ and NGF^Wild^, respectively, where a positive percentage represents that NGF^R100W^ induced the bond frequency, and a negative percentage denotes that NGF^R100W^ reduced the bond frequency.

## 5 CONCLUSION

Elucidating the mechanism of developing HSAN V disorder, characterized by insensitivity to pain due to NGF^R100W^, is indispensable to understanding how altered dynamics motion in NGF^R100W^ abolishes NGF-TrkA-triggered signaling for pain sensation and induced pathophysiology of the HSAN V disease. Through an extensive MD simulation study, this study discloses the mechanism of how NGF^R100W^ altered the dynamics motion of TrkA particulate for pain sensation. Findings from this study can be used to apprehend the pathophysiology of HSAN V disorder as well as to develop novel therapeutics that eliminate the differential binding profiles of NGF-TrkA, so restoring the native conformational dynamics, motions, and direction of structural movements of these residues might be an appealing approach for addressing NGF^R100W^-induced HSAN V disorder.

## Author Contributions

The authors certify their contributions to this work as follows: Conceptualization, R.D.; methodology, Z.B.R., M.A.B., and N.D.; software, Z.B.R., M.A.B., N.D.; validation, Z.B.R., M.A.B., N.D., S.P., R.Z., and S.M.; formal analysis, Z.B.R., M.A.B., N.D., S.P., R.Z.; investigation, Z.B.R., M.A.B., N.D., S.P., R.Z., S.M., and R.D.; resources, R.D.; visualization, Z.B.R., M.A.B., S.P., and R.D.; writing–original draft preparation, M.A.B., and R.D.; writing–review and editing, M.A.B., S.M., and R.D.; project administration, R.D. Every author has reviewed and approved the finalized draft of the manuscript. Z.B.R.^#^ and M.A.B.^#^ equally contributed to this work.

## Notes

The authors declare that there are no competing financial or any other interests.

