## Supplementary File for "Unraveling the distinct motion bias of TrkA-NGF complex in NGF^R100W^-driven HSAN V disease"

**Table S1.** Total MM-GBSA binding free energy ($\Delta G_{bind}$) between $NGF_V$ and $NGF_W$ Chain of NGF dimer.

| Clusters | Wild (kcal/mol) | R100W (kcal/mol) |
|---|---|---|
| Cluster 1 | -178.53 | -156.68 |
| Cluster 2 | -177.8 | -156.54 |
| Cluster 3 | -171.81 | -160.17 |
| Cluster 4 | -177.2 | -155.76 |
| Cluster 5 | -177.62 | -151.9 |
| Cluster 6 | -171.72 | -152.85 |
| Cluster 7 | -190.48 | -162.13 |
| Cluster 8 | -178.43 | -161.55 |
| Cluster 9 | -191.96 | -151.55 |
| Cluster 10 | -178.33 | -157.01 |
| **Mean ± SD** | -179.39 ± 6.75 | -156.61 ± 3.81 |

**Table S2.** Total MM-GBSA binding free energy ($\Delta G_{bind}$) between NGF dimer and $TrkA_X$ and $TrkA_Y$ chain.

| Clusters | $TrkA_X$ Chain | | $TrkA_Y$ Chain | |
|---|---|---|---|---|
| | Wild (kcal/mol) | R100W (kcal/mol) | Wild (kcal/mol) | R100W (kcal/mol) |
| Cluster 1 | -73.22 | -103.89 | -70.65 | -102.03 |
| Cluster 2 | -80.65 | -103.87 | -73.54 | -98.19 |
| Cluster 3 | -81.76 | -103.73 | -77.59 | -97.76 |
| Cluster 4 | -87.24 | -102.73 | -78.16 | -97.04 |
| Cluster 5 | -89.17 | -102.47 | -78.25 | -96.64 |
| Cluster 6 | -89.35 | -101.36 | -79.24 | -96.55 |
| Cluster 7 | -89.76 | -99.31 | -79.94 | -96.36 |
| Cluster 8 | -90.17 | -98.32 | -82.47 | -92.1 |
| Cluster 9 | -91.84 | -98.32 | -84.59 | -91.66 |
| Cluster 10 | -93.11 | -97.52 | -85.18 | -90.7 |
| **Mean ± SD** | -86.63 ± 6.19 | -101.15 ± 2.55 | -78.96 ± 4.54 | -95.90 ± 3.47 |

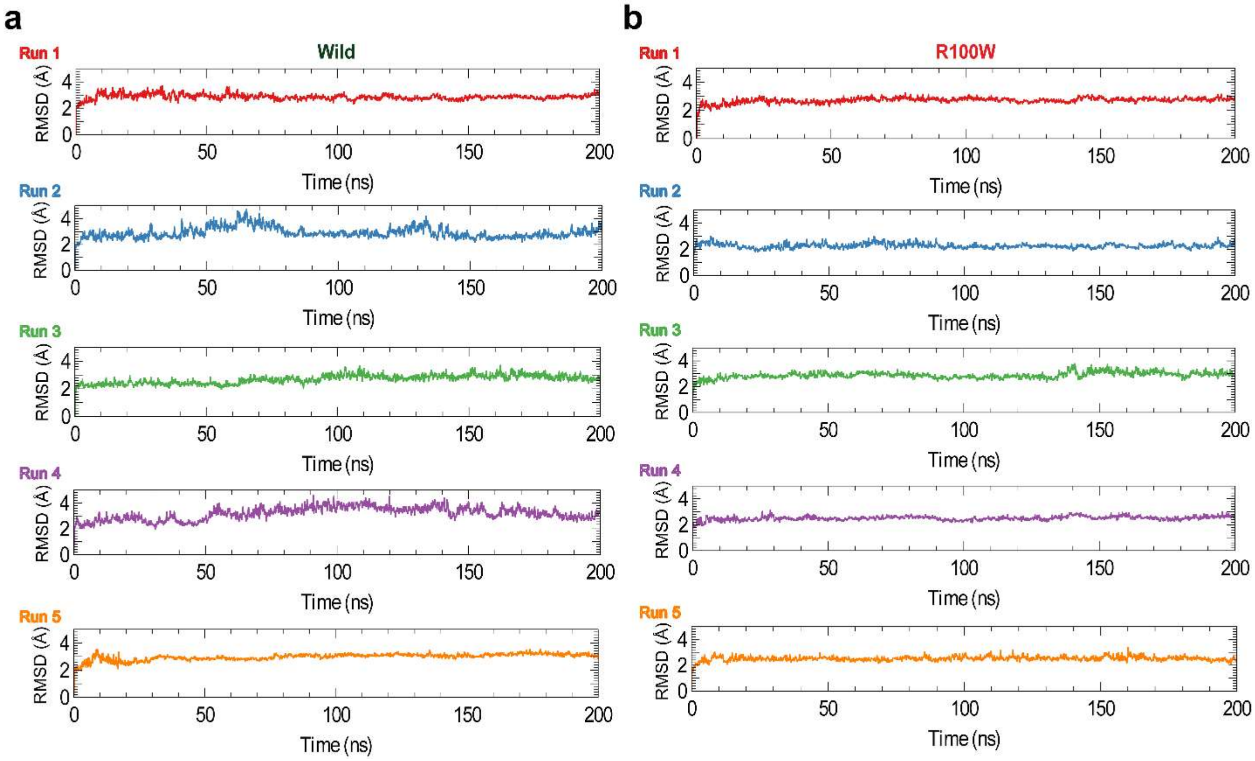


**Figure S1.** The RMSD values obtained from each run of different conditions for wild and mutant-containing structures were evaluated using protein C-alpha of (a) $NGF^{Wild}$ and (b) $NGF^{R100W}$, respectively.

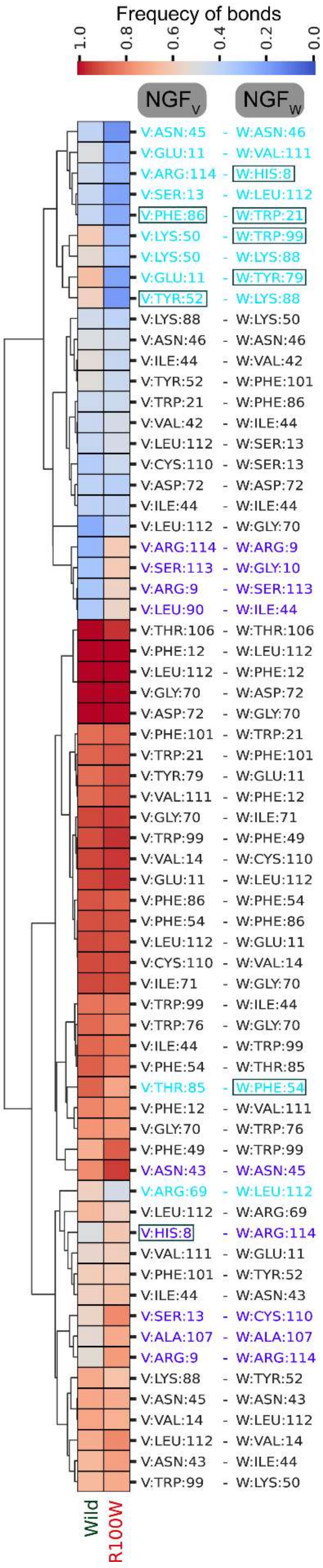

Frequency of bonds
1.0
0.8
0.6
0.4
0.2
0.0
NGF$_V$
NGF$_W$
V:ASN:45 - W:ASN:46
V:GLU:11 - W:VAL:111
V:ARG:114 - W:HIS:8
V:SER:13 - W:LEU:112
V:PHE:86 - W:TRP:21
V:LYS:50 - W:TRP:99
V:LYS:50 - W:LYS:88
V:GLU:11 - W:TYR:79
V:TYR:52 - W:LYS:88
V:LYS:88 - W:LYS:50
V:ASN:46 - W:ASN:46
V:ILE:44 - W:VAL:42
V:TYR:52 - W:PHE:101
V:TRP:21 - W:PHE:86
V:VAL:42 - W:ILE:44
V:LEU:112 - W:SER:13
V:CYS:110 - W:SER:13
V:ASP:72 - W:ASP:72
V:ILE:44 - W:ILE:44
V:LEU:112 - W:GLY:70
V:ARG:114 - W:ARG:9
V:SER:113 - W:GLY:10
V:ARG:9 - W:SER:113
V:LEU:90 - W:ILE:44
V:THR:106 - W:THR:106
V:PHE:12 - W:LEU:112
V:LEU:112 - W:PHE:12
V:GLY:70 - W:ASP:72
V:ASP:72 - W:GLY:70
V:PHE:101 - W:TRP:21
V:TRP:21 - W:PHE:101
V:TYR:79 - W:GLU:11
V:VAL:111 - W:PHE:12
V:GLY:70 - W:ILE:71
V:TRP:99 - W:PHE:49
V:VAL:14 - W:CYS:110
V:GLU:11 - W:LEU:112
V:PHE:86 - W:PHE:54
V:PHE:54 - W:PHE:86
V:LEU:112 - W:GLU:11
V:CYS:110 - W:VAL:14
V:ILE:71 - W:GLY:70
V:TRP:99 - W:ILE:44
V:TRP:76 - W:GLY:70
V:ILE:44 - W:TRP:99
V:PHE:54 - W:THR:85
V:THR:85 - W:PHE:54
V:PHE:12 - W:VAL:111
V:GLY:70 - W:TRP:76
V:PHE:49 - W:TRP:99
V:ASN:43 - W:ASN:45
V:ARG:69 - W:LEU:112
V:LEU:112 - W:ARG:69
V:HIS:8 - W:ARG:114
V:VAL:111 - W:GLU:11
V:PHE:101 - W:TYR:52
V:ILE:44 - W:ASN:43
V:SER:13 - W:CYS:110
V:ALA:107 - W:ALA:107
V:ARG:9 - W:ARG:114
V:LYS:88 - W:TYR:52
V:ASN:45 - W:ASN:43
V:VAL:14 - W:LEU:112
V:LEU:112 - W:VAL:14
V:ASN:43 - W:ILE:44
V:TRP:99 - W:LYS:50
Wild
R100W

**Figure S2.** The illustration demonstrates total contact alteration by $NGF^{Wild}$ and $NGF^{R100W}$ between $NGF_V$ and $NGF_W$ chains, where the heatmap describes the frequency of total bonds between $NGF_V$ and $NGF_W$ chains. As shown in the frequency heatmap due to $NGF^{R100W}$, reduced bonds are shown in light blue, while enhanced bonds are shown in violet. Aromatic residues whose bond frequency has changed by $NGF^{R100W}$ are marked with a box.

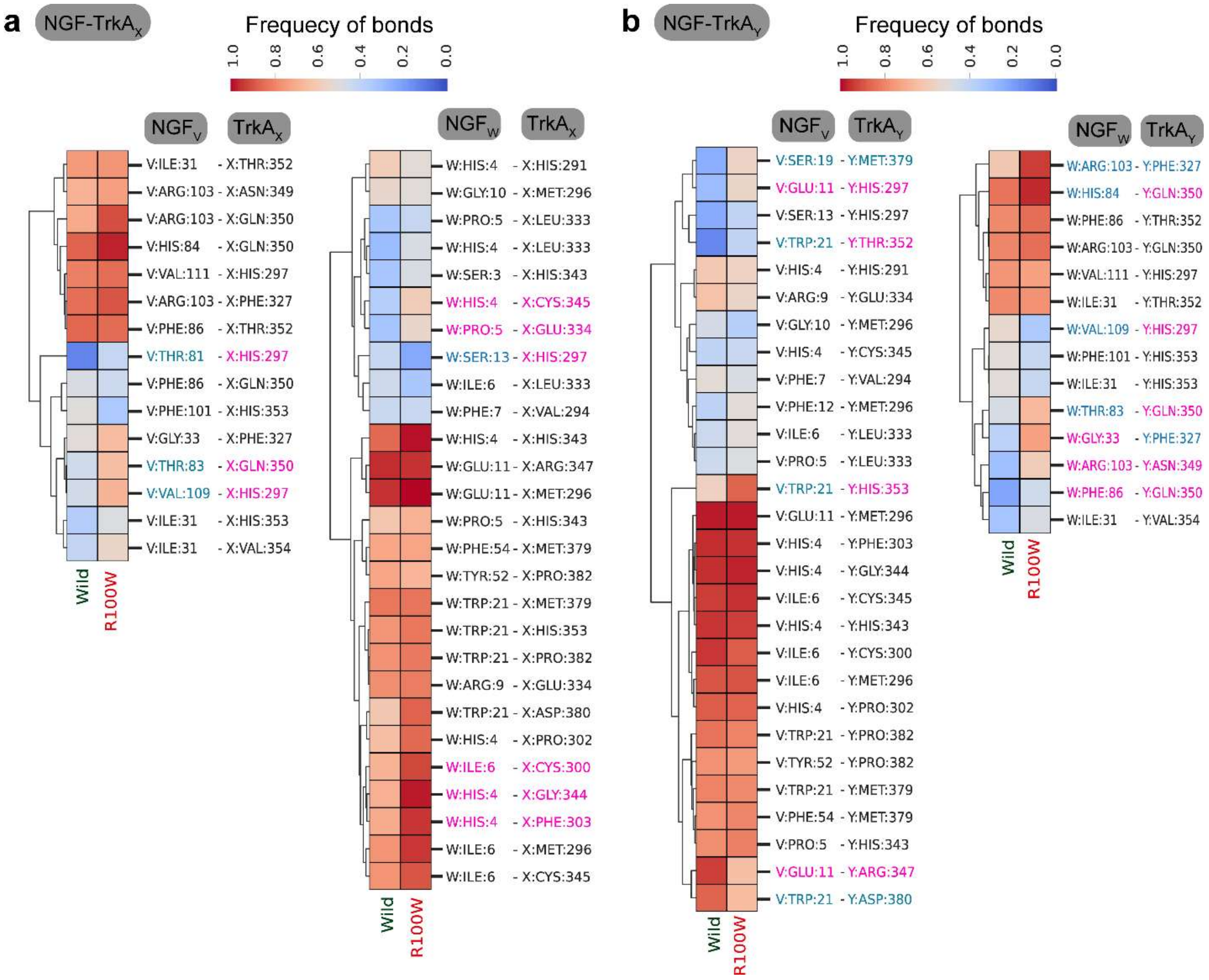


**Figure S3.** The graphical representation demonstrates total contact alteration between NGF and TrkA$_X$ chain (a) and NGF and TrkA$_Y$ chain (b) due to NGF$^{Wild}$ and NGF$^{R100W}$, where the heatmap describes the frequency of total bonds between NGF and TrkA with red color representing frequency 1. The residues from the specificity patches and common patches are shown by the colors pink and blue, respectively.

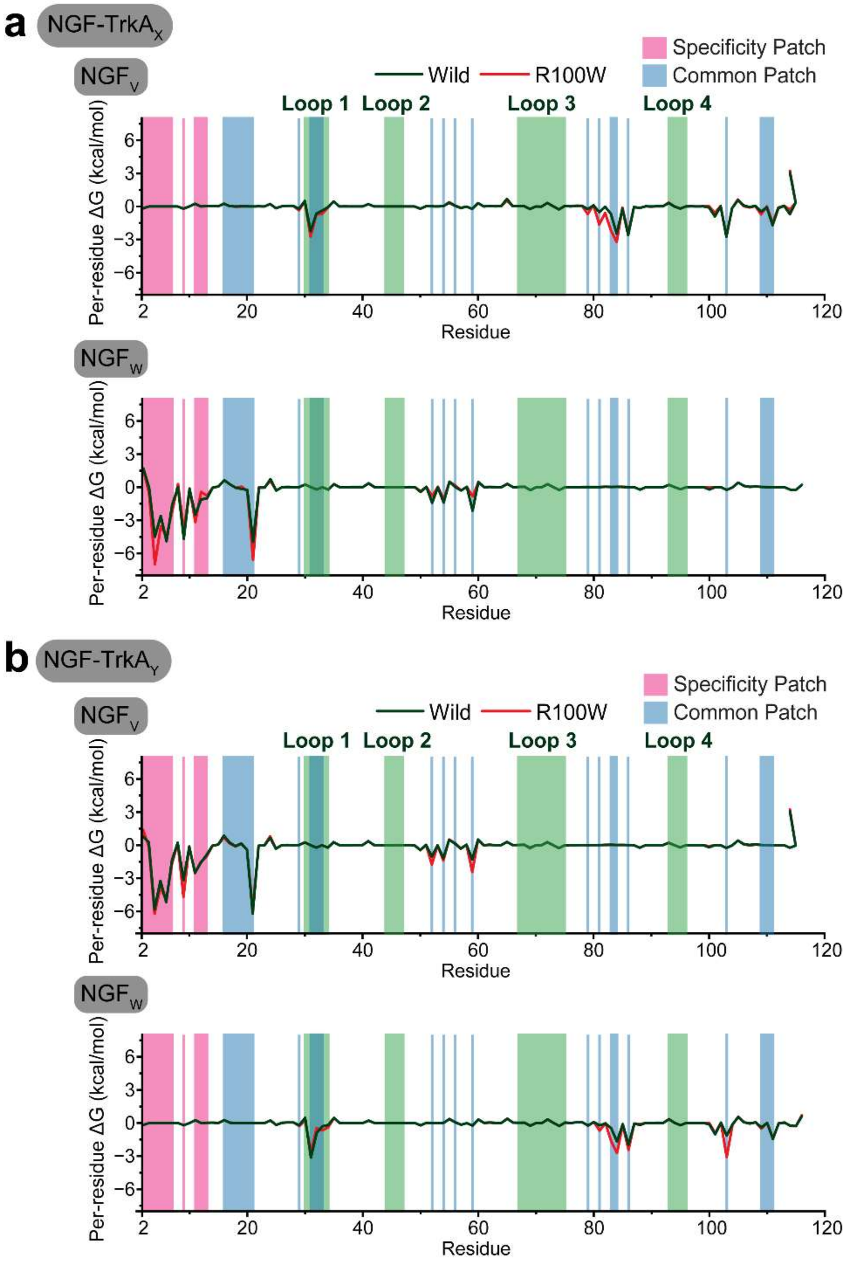


**Figure S4.** The per-residue decomposition analyses demonstrate the binding contribution of (a) NGF$_V$ (upper panel) and NGF$_W$ (lower panel) with TrkA$_X$ and (a) NGF$_V$ (upper panel) and NGF$_W$ (lower panel) with TrkA$_Y$ between NGF$^{Wild}$ and NGF$^{R100W}$. The loop regions (1, 2, 3, and 4) are represented by the color green, while the specificity patches and common patches are shown by the colors pink and blue, respectively.

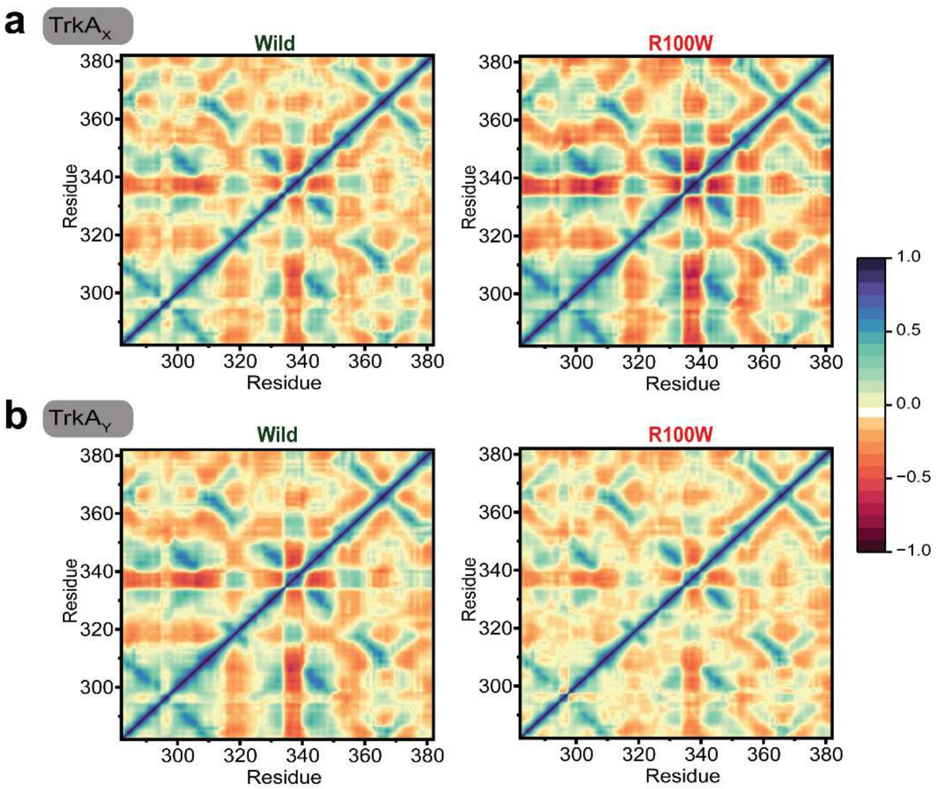


**Figure S5.** The dynamic cross-correlation matrix (DCCM) highlights the anticorrelated and correlated motions of each residue pair in the structures, $NGF^{Wild}$ and $NGF^{R100W}$ for $TrkA_X$ chain (a) and $TrkA_Y$ chain (b). A highly correlated motion is symbolized by red (+1), while an anticorrelated motion is represented by blue (−1).

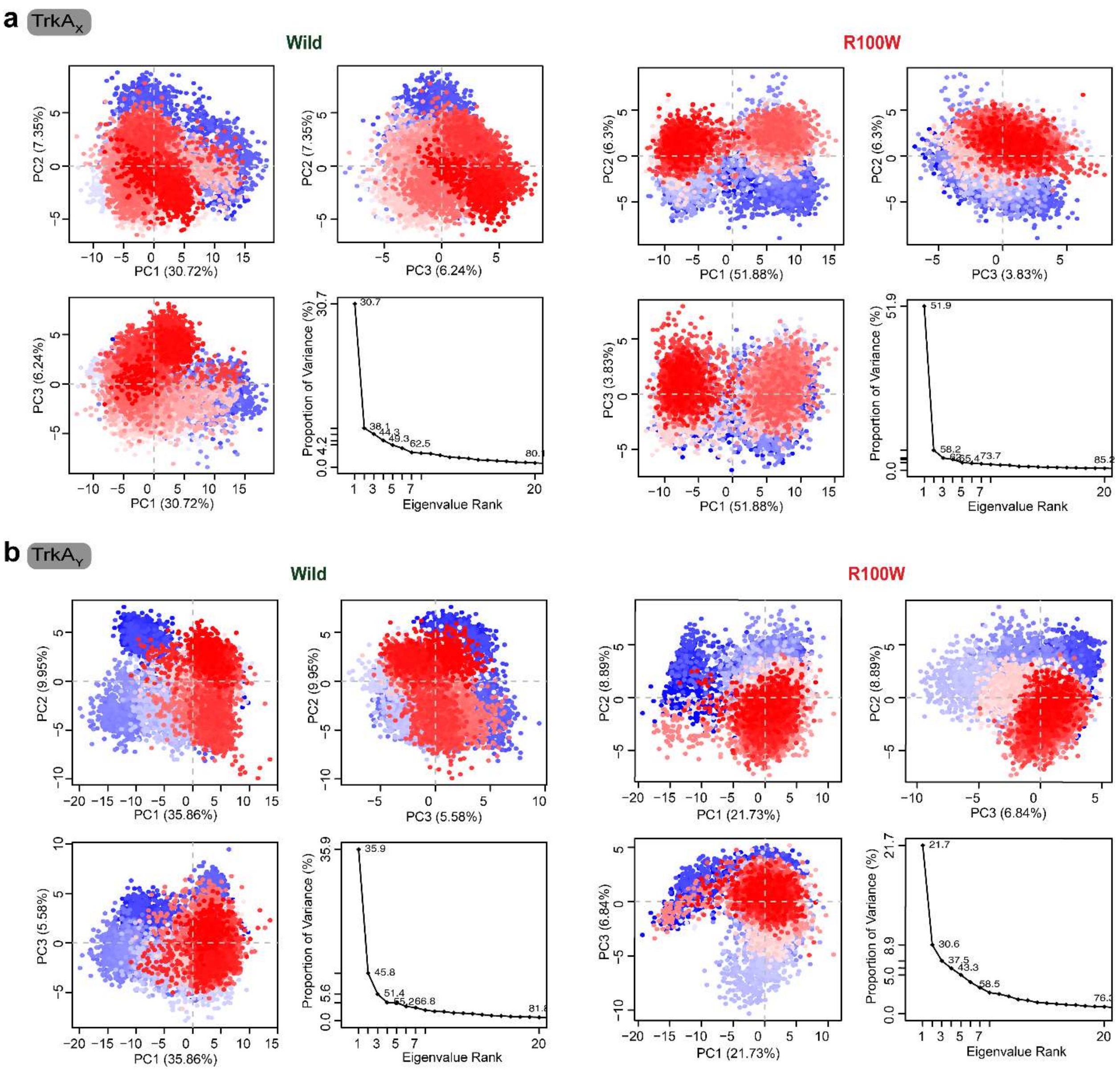


**Figure S6.** The trajectory conformer distribution was displayed over the principal planes components (PC) 1, 2 and 3, with each dot symbolizing a structure that changed color with time (from blue to red). The first 3 plots demonstrate comparison among PC1, PC2 and PC3 of NGF$^{Wild}$ and NGF$^{R100W}$ in the TrkA$_X$ chain (a) and TrkA$_Y$ chain (b). The fourth plot represents the proportion of variance for each PC of NGF$^{Wild}$ and NGF$^{R100W}$ in the TrkA$_X$ (a) and TrkA$_Y$ chain (b).

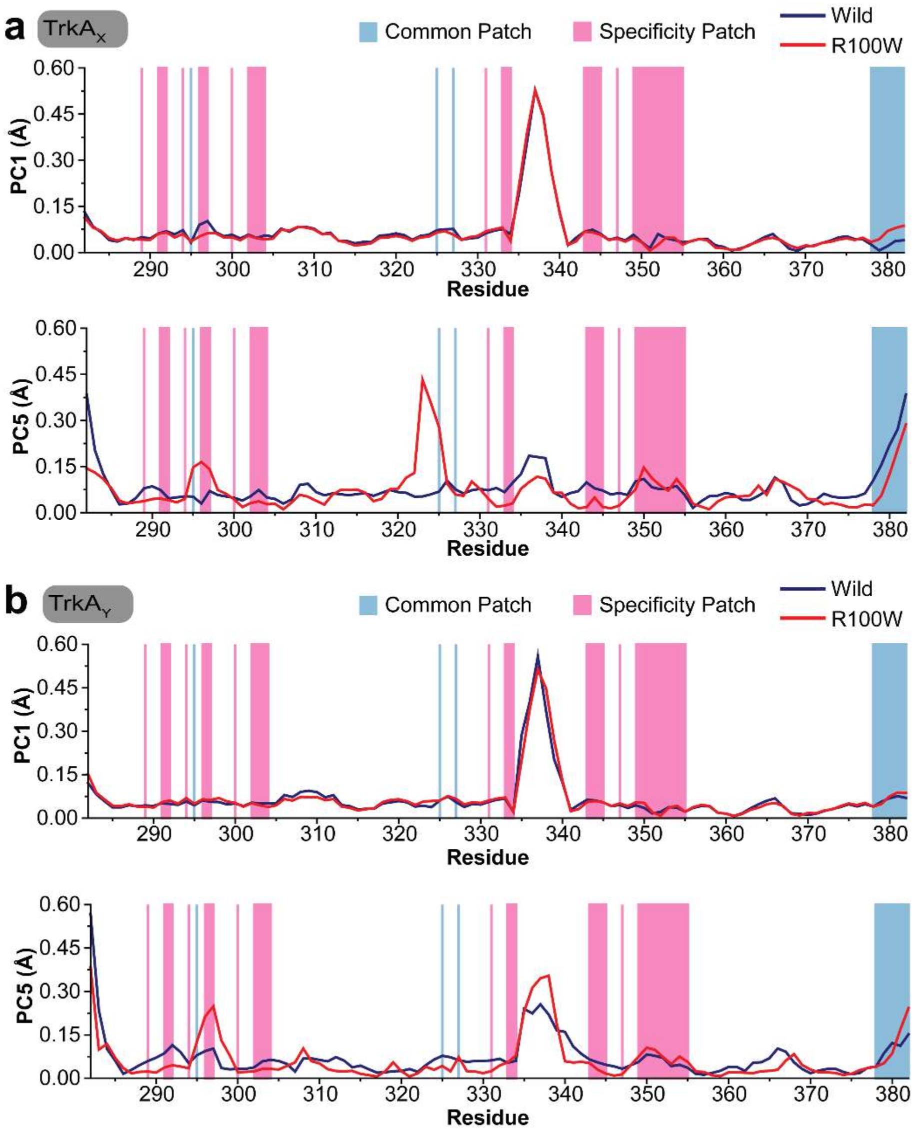


**Figure S7.** The graphical representations display line plots illustrating the mobility degree captured by PC1 and PC5 for the TrkA$_X$ chain (a) and TrkA$_Y$ chain (b). The specificity patches and common patches are shown by the colors pink and blue, respectively.
